# Safety Transparency in Animal Cell-Cultured Ingredients for Pet Food: A Case Study Establishing the Standard for Public Disclosure

**DOI:** 10.64898/2026.07.14.738473

**Authors:** Rupal Tewari, Robert Soukup, Loukia Hadjistylianou, Marta Manicone, Marta Serra, Manuel Felbermair, Shannon Falconer

## Abstract

Animal cell-cultured ingredients are entering the EU and UK pet food markets under frameworks that do not require pre-market, ingredient-level safety assessments, creating an ethical need for transparent safety disclosure. We present the first public safety dossier for this sector, describing the proprietary mouse embryonic stem cell line PE25 and its derived, non-viable cellular and conditioned media ingredient produced in food and feed-grade media. PE25 characterization confirmed *Mus musculus* identity, sterility, absence of mycoplasma and replication-competent retroviruses, and stable growth. Doxorubicin-induced p53 stress testing, *CD44/BMI1* profiling, and soft agar assays showed no cancer-like traits and a non-tumorigenic profile; the final ingredient contains no viable cells. Independent OECD TG 471 and 487 assays confirmed non-genotoxicity. Heavy metals, biogenic amines, solvents, and chemical residues were below regulatory limits. Given process variability, we recommend case-by-case safety evaluation and propose this dossier as a model for responsible commercialization.

## Introduction

The commercial development of animal cell-cultured ingredients for pet food has attracted considerable attention, partly because the regulatory pathway for animal feed is generally perceived to carry a lower barrier to entry than that for human food. Within the European Union, feed materials are governed by EU Feed Regulation (EC) No 767/2009, under which a manufacturer may self-register a new feed material on the EU Feed Materials Register without first submitting a safety dossier to the European Food Safety Authority (EFSA) or any other independent authority for scientific review^1^.

The obligation placed on the manufacturer is accuracy of labelling and general compliance with feed safety principles—not the evidence-based pre-market authorization required under the EU Novel Food Regulation (EU) No 2015/2283, which mandates a full EFSA safety opinion before market entry^2,3^.

A cell-cultured ingredient entering the EU pet food market via the feed materials route is therefore not subject to independent third-party vetting of its specific safety profile prior to commercialization^4,5^. In the UK, the Food Standards Agency’s Cell-Cultivated Products (CCP) sandbox is actively developing a regulatory pathway for cultivated products and aims to complete risk assessments for initial CCP applications by February 2027^6^. In the interim, no pre-market authorization requirement equivalent to that of the EU Novel Food Regulation currently applies to cell-cultured ingredients destined for pet food in the UK. These regulatory realities are not a criticism of the frameworks themselves—both are in active and constructive development—but they do create a situation in which companies can lawfully bring animal cell-cultured pet food ingredients to market without publicly disclosing comprehensive ingredient-level safety data. Rather than treating this lower regulatory bar as an opportunity, manufacturers have an ethical obligation to treat it as a responsibility. The absence of mandatory independent third-party vetting should not translate into the absence of independent third-party safety testing. The precedent set here—in the earliest days of this industry—will define public trust in animal cell-cultured ingredients for years to come.

A further challenge is the tendency to treat “animal cell-cultured ingredients” as a monolithic category in which the safety assessment of one product speaks for all. This is scientifically untenable. Cell-cultured ingredients can differ substantially in cell line species and tissue of origin, culture media composition, serum-free adaptation status, passage history, processing aids employed, and the scope of safety testing performed^7,8^. An ingredient produced from an immortalized cell line in undefined animal-derived media after hundreds of passages presents a fundamentally different safety profile from one derived from embryonic stem cells in food-grade media and subjected to comprehensive multi-tier safety characterization. Regulators and pet food manufacturers must evaluate each product individually, and manufacturers must proactively supply the data to make that evaluation possible^9^.

One of the most prominent public concerns regarding animal cell-cultured ingredients is whether the cell lines used to produce them exhibit tumorigenic or cancer-like properties. While the scientific consensus holds that consumption of tumorigenic cells does not constitute a direct health hazard—the digestive process degrades cells into their constituent molecular components—consumer and regulatory expectations demand that ingredients intended for food and feed use are derived from cells that do not display the hallmarks of malignant transformation^10^. Demonstrating the absence of such properties is therefore an important component of responsible product characterization, independent of any direct toxicological risk.

The cell biology underlying this concern is well-documented and spans multiple cell types^11^. For deliberately immortalized cell lines—among the most commonly used in the cultivated meat industry—the mechanism of immortalization is itself a key variable. Strategies based on direct inactivation of the tumor protein (p53) and retinoblastoma protein (Rb1) tumour suppressor pathways functionally disable the cell’s primary genomic surveillance checkpoints^12,13^. Approaches based on telomerase reverse transcriptase (TERT) overexpression are generally considered less disruptive, though co-expression with cell cycle accelerators such as cyclin dependent kinase 4 (CDK4) introduces partial checkpoint bypass^14^. Spontaneous immortalization—in which cells acquire indefinite proliferative capacity without deliberate genetic engineering, through endogenous mechanisms such as telomerase activation—is frequently presented as a preferable alternative^15,16^, but the integrity of tumour suppressor pathways in spontaneously immortalized lines cannot be assumed without direct empirical testing^17^. For pluripotent stem cell lines including embryonic stem cells, chromosomal instability and recurrent deletions at the *TP53* locus have been documented under specific long-term culture conditions, particularly with enzymatic passaging in feeder-free systems^18^.

Across all of these cell types, the relevant question is whether the cell line, as actually used in production, retains functional tumour suppressor activity and lacks malignant phenotypic properties. This cannot be answered by characterizing the immortalization method alone, nor resolved by theoretical arguments about cell type. It requires direct empirical data, reported transparently, for each individual animal cell line.

A rigorous safety framework for a cell-cultured ingredient begins with the cell bank itself. A tiered cell banking system—comprising a Parental Cell Bank (PCB), a Master Cell Bank (MCB), and a Working Cell Bank (WCB) established in accordance with International Council for Harmonisation of Technical Requirements for Registration of Pharmaceuticals for Human Use-Quality 5D (ICH Q5D) guidelines provides the foundational traceability and biological consistency upon which all downstream safety claims rest^19^. Without a documented, characterized cell bank, passage numbers are unverifiable, lot-to-lot consistency cannot be assured, and the basis for safety claims is undermined. Cell bank characterization must include confirmation of species identity, demonstration of sterility and absence of mycoplasma, and assessment for adventitious viral agents. This last point warrants particular attention for rodent-derived cell lines: mouse and other murine cell lines harbour endogenous retroviral elements (ERVs) integrated into the host genome, which represent a characteristic of murine genomic biology rather than an acquired contamination^20,21^. The critical safety question is not whether such sequences are present—they are expected and their presence in next generation sequencing (NGS) data does not constitute a safety concern—but whether replication-competent, infectious viral particles capable of transmission are absent. Demonstrating the absence of replicative infectivity requires a functional transmissibility assay, not transcriptomic detection alone^22,23^.

Beyond cell-level characterization, a responsible safety dossier must address the chemical profile of the final ingredient as it reaches the consumer’s bowl. Bioreactor-based production introduces potential contamination routes—from equipment surfaces, media components, and processing reagents—that have no equivalent in conventional food manufacturing^24^. The use of dimethyl sulfoxide (DMSO) as a cryoprotectant in cell banking is standard practice^25^, but its controlled removal before the production seed train, and its confirmed absence in the harvested ingredient, must be documented. Biogenic amines can accumulate in protein-rich biological matrices under conditions of microbial activity or cellular degradation^26^; although certain biogenic amines are known to possess positive functional benefits^27^, others are relevant when considering ingredient safety^28,29^. Heavy metals and volatile organic solvents must be screened against applicable ICH Q3C (residual solvents scientific guideline) using validated, accredited analytical methods^30^. Taken together, these screens constitute the chemical safety tier of a complete ingredient dossier.

Media transparency is equally important. Without public disclosure of what a cell-cultured ingredient was grown in, neither regulators nor manufacturers downstream in the supply chain can meaningfully assess contamination risk, evaluate residue testing completeness, or make informed procurement decisions. The regulatory approval status of every media component under applicable feed legislation should be considered a minimum disclosure requirement.

For any ingredient derived from a novel production system, genotoxicity testing of the finished product—rather than its individual components—provides the most direct and informative safety assurance that the final material is neither mutagenic, clastogenic nor aneugenic^31^. The Ames bacterial reverse mutation assay^32^ (OECD TG 471) and the in vitro mammalian cell micronucleus test^33^ (OECD TG 487) represent the internationally harmonized minimum standard for food-relevant substance genotoxicity screening, as recommended by EFSA^34^, and the FDA^35^. Together, the Ames bacterial reverse mutation assay and the in vitro mammalian cell micronuclease assay cover all three principal genotoxic endpoints: gene mutation, structural chromosomal aberration (clastogenicity), and numerical chromosomal aberration (aneugenicity). For a cell-cultured ingredient whose production media contains a defined formulation of amino acids, vitamins, minerals, and metabolic substrates—whose individual components are approved, but whose interactions in a complex biological matrix have not been independently evaluated—testing the harvested ingredient as a whole is the only approach that addresses real-world exposure.

This paper presents a comprehensive safety characterization of proprietary mouse embryonic stem cell (mESC) line PE25 and its derived animal cell-cultured ingredient—a combination of non-viable harvested cells and conditioned media, produced entirely in food-safe and feed-grade media, complying with the EU Directive 2002/32/EC on undesirable substances in animal feed^36^; EU directive 2009/32/EC on extraction solvents used in the production of foodstuffs and the food ingredients^37^; EU commission regulation (EU) No. 68/2013 on the catalogue of feed materials^38^; and EU commission regulation (EC) No 1831/2003 on additives for use in animal nutrition^39^, all components of which hold current EU regulatory approval (**Table 1**). To our knowledge, this constitutes the first public disclosure of a complete cell line and ingredient safety dossier by any company producing animal cell-cultured ingredients for pet food. The aims of this work are: (1) to demonstrate that rigorous, multi-tier safety testing is achievable at the ingredient-level; (2) to establish, using three independent assays, that cell line PE25 is non-tumorigenic and does not display cancer-associated properties; (3) to present cell bank characterization data including viral safety assessment for a murine cell line; (4) to report chemical safety data—including heavy metals, solvent residuals, and biogenic amines—for the harvested ingredient using accredited third-party analytical methods; (5) to present independent third-party genotoxicity data (OECD TG 471 and TG 487) representing an unprecedented level of safety scrutiny for a commercial animal cell-cultured pet food ingredient; and (6) to propose a minimum safety disclosure standard that we call on the broader industry to adopt.

**Table 1.**
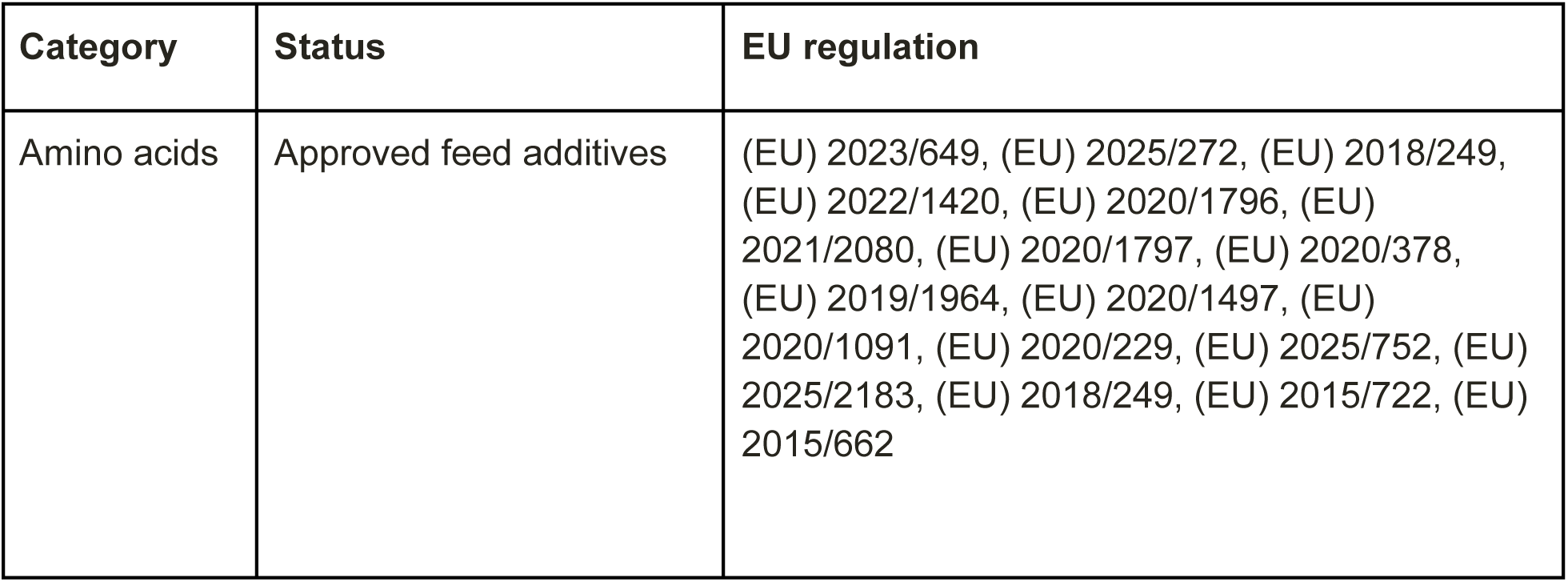
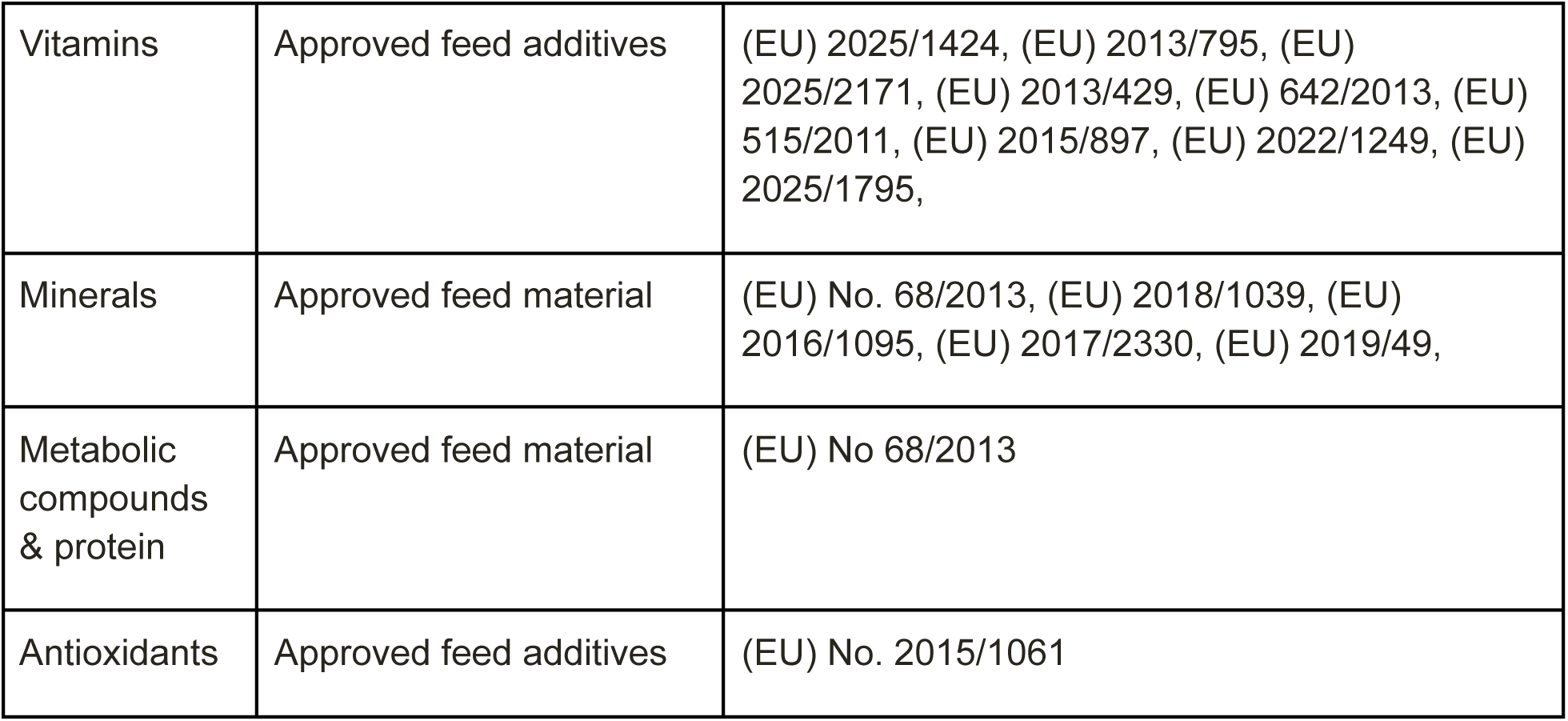
Regulatory approval status of production media components under EU feed legislation.

This paper is intended to serve as both a scientific record and a practical template for others in the industry to follow.

## Results

### MCB Characterization: Identity, Sterility, and Viral Safety

Species identity testing by quantitative polymerase chain reaction (qPCR) confirmed the PE25 cell line used to generate the MCB as *Mus musculus* with no evidence of cross-contamination. Sterility testing by direct inoculation per European Pharmacopoeia (EP 2.6.1) / US Pharmacopoeia (USP 71) returned negative results, and mycoplasma/Spiroplasma testing by qPCR per EP 2.6.7 / USP 63 likewise returned negative results. NGS transcriptomic analysis detected genomic sequences of Murine Leukemia Virus (MLV) in the PE25 MCB cells; these represent ERVs integrated into the mouse genome through evolutionary processes and are distinct from exogenous, replication-competent particles^40^. To confirm the absence of infectious particles, a transmissibility assay was performed by co-culturing MCB cells with *Mus dunni* target cells using a quantitative product enhanced reverse transcriptase (Q-PERT) readout. Results were negative for replicative retrovirus infectivity.

### WCB: Adaptation to Food-Safe and Feed-Grade Media

The PE25 WCB was successfully established after 46 passages in 100% food-safe and feed-grade media (**Table 1**); all components hold current approval as EU Feed Additives or are listed in Feed Materials catalogue. Establishing the WCB involved a selection process allowing for clonal adaptation to the food-safe and feed-grade formulation.

### Chemical Safety: Heavy Metals, Solvents, and Biogenic Amines

Inductively coupled plasma mass spectrometry (ICP-MS) analysis confirmed the absence of lead (Pb), cadmium (Cd), arsenic (As), and mercury (Hg) above the limits of quantification in both the PE25 WCB cell biomass and conditioned media (**Table 2**). Of the 16 volatile organic solvents screened (benzene, methanol, ethanol, acetone, isopropanol, toluene, ethylbenzene, *m,p*-xylene, *o*-xylene, chloroform, dichloromethane, trichloroethylene, 1,1,1-trichloroethane, tetrachloroethylene, carbon tetrachloride and DMSO), with the exception of ethanol, all were below the respective limits of quantification (**Table 2**). β-mercaptoethanol (BME)—used as a processing aid prior to establishment of the WCB—was also not detected. Of the nine biogenic amines screened, (histamine, cadaverine, putrescine, spermidine, spermine, tyramine, phenylethylamine, tryptamine, and isopentylamine), with the exception of spermine, all were below the limit of quantification (**Table 2**).

**Table 2.** Heavy metal, solvent and chemical reagent residue, and biogenic amine screening of PE25 WCB cells and conditioned media.

| Parameter | Method | Unit | Result | Limit of Quantification |
| --- | --- | --- | --- | --- |
| **Heavy metals** |  |  |  |  |
| Lead (Pb) | DIN EN 15763, mod. | mg/kg | < 0.015 | 0.015 |
| Cadmium (Cd) | DIN EN 15763, mod. | mg/kg | < 0.010 | 0.010 |
| Mercury (Hg) | DIN EN 15763, mod. | mg/kg | < 0.010 | 0.010 |
| Arsenic (As) | DIN EN 15763, mod. | mg/kg | < 0.04 | 0.04 |
| **Solvents** |  |  |  |  |
| Benzene | HS-GC/MS, SOP M 1299 | mg/kg | < 0.030 | 0.030 |
| Toluene | HS-GC/MS, SOP M 1299 | mg/kg | < 0.030 | 0.030 |
| Methanol | HS-GC-FID | mg/kg | < 10 | 10 |
| Ethanol | HS-GC-FID | mg/kg | 6 | 1 |
| Acetone | HS-GC-FID | mg/kg | < 1 | 1 |
| Isopropanol | HS-GC-FID | mg/kg | < 1 | 1 |
| Ethylbenzene | HS-GC/MS, SOP M 1299 | mg/kg | < 0.030 | 0.030 |
| m,p-xylene | HS-GC/MS, SOP M 1299 | mg/kg | < 0.030 | 0.030 |
| o-xylene | HS-GC/MS, SOP M 1299 | mg/kg | < 0.030 | 0.030 |
| Chloroform | HS-GC/MS, SOP M 1299 | mg/kg | < 0.010 | 0.010 |
| Dichloromethane | HS-GC/MS, SOP M 1299 | mg/kg | < 0.010 | 0.010 |
| Trichloroethylene | HS-GC/MS, SOP M 1299 | mg/kg | < 0.010 | 0.010 |
| 1,1,1-trichloroethane | HS-GC/MS, SOP M 1299 | mg/kg | < 0.010 | 0.010 |
| Tetrachloroethylene | HS-GC/MS, SOP M 1299 | mg/kg | < 0.010 | 0.010 |
| Carbon tetrachloride | HS-GC/MS, SOP M 1299 | mg/kg | < 0.010 | 0.010 |
| DMSO | GC-MS | mg/kg | < 1 | 1 |
| **Biogenic amines** |  |  |  |  |
| Histamine | HPLC, SOP M 1574 | mg/kg | < 50 | 50 |
| Cadaverine | HPLC, SOP M 1574 | mg/kg | < 50 | 50 |
| Putrescine | HPLC, SOP M 1574 | mg/kg | < 50 | 50 |
| Spermidine | HPLC, SOP M 1574 | mg/kg | < 50 | 50 |
| Spermine | HPLC, SOP M 1574 | mg/kg | 71 | 50 |
| Tyramine | HPLC, SOP M 1574 | mg/kg | < 50 | 50 |
| Phenylethylamine | HPLC, SOP M 1574 | mg/kg | < 50 | 50 |
| Tryptamine | HPLC, SOP M 1574 | mg/kg | < 50 | 50 |
| Isopentylamine | HPLC, SOP M 1574 | mg/kg | < 50 | 50 |
| **Chemical reagent** |  |  |  |  |
| BME | GC-MS | mg/kg | < 0.03 | 0.1 |

To ensure complete removal of DMSO carried over from WCB cryovials, a three-step clearance protocol was employed. First, cells were subjected to an immediate 10-fold dilution upon thawing, reducing the extracellular DMSO concentration from 10% (110 mg/mL) to 1% (11 mg/mL). During post-thaw dilution, the extracellular DMSO concentration is rapidly reduced as intracellular DMSO remains temporarily elevated. This creates an outward concentration gradient that drives passive efflux of DMSO from the cell^41, 42, 43^. Second, a double-wash step eliminated the remaining extracellular DMSO, sustaining the outward gradient and enabling continued passive efflux. Third, cells underwent a minimum of eight passages at a 1:5 split ratio in DMSO-free, food-safe and feed-grade production media prior to entering the seed train; successive cell divisions further diluted any residual intracellular DMSO content across daughter cells.

Third-party gas chromatography-mass spectrometry (GC-MS) quantification of DMSO in PE25 WCB cells and conditioned media at passage 8 post-thaw confirmed DMSO below the limit of quantification (LOQ: 1 mg/kg), the lowest LOQ any consulted third-party laboratory was able to achieve. Under the most conservative theoretical assumptions—accounting for the initial 10-fold dilution, double-wash, and eight passages of 1:5 serial dilution (a mathematical 390,625-fold dilution)—the maximum residual DMSO concentration after the three-step process is below 2.82 x 10⁻^5^ mg/mL.

For context, landmark chronic toxicity data establish that the threshold for DMSO toxic effects is approximately 1,100 mg/kg body weight/day in dogs and 9,900 mg/kg body weight/day in rats^44^. Assuming a conservative dietary inclusion rate of 50% of PE25 plus conditioned media, and a standard daily food intake of 2.5% of body weight, a dog consuming this ingredient in which DMSO remained at the LOQ (1 mg/kg) would be exposed to approximately 0.0125 mg DMSO/kg body weight/day—a safety margin exceeding 88,000-fold below the established toxicity threshold.

### Passage Stability: Consistent Doubling Time Across the Production Passage Range

Doubling time of PE25 WCB cells remained consistent throughout the monitored passage range, with no systematic trend towards acceleration or deceleration that would indicate progressive genomic destabilization or crisis (**Fig. 1**). The mean doubling time was 22.8 hours. This stability is consistent with a cell population that has reached a stable adapted state rather than one undergoing ongoing selective genomic disruption^45,46^.

**Fig. 1.**
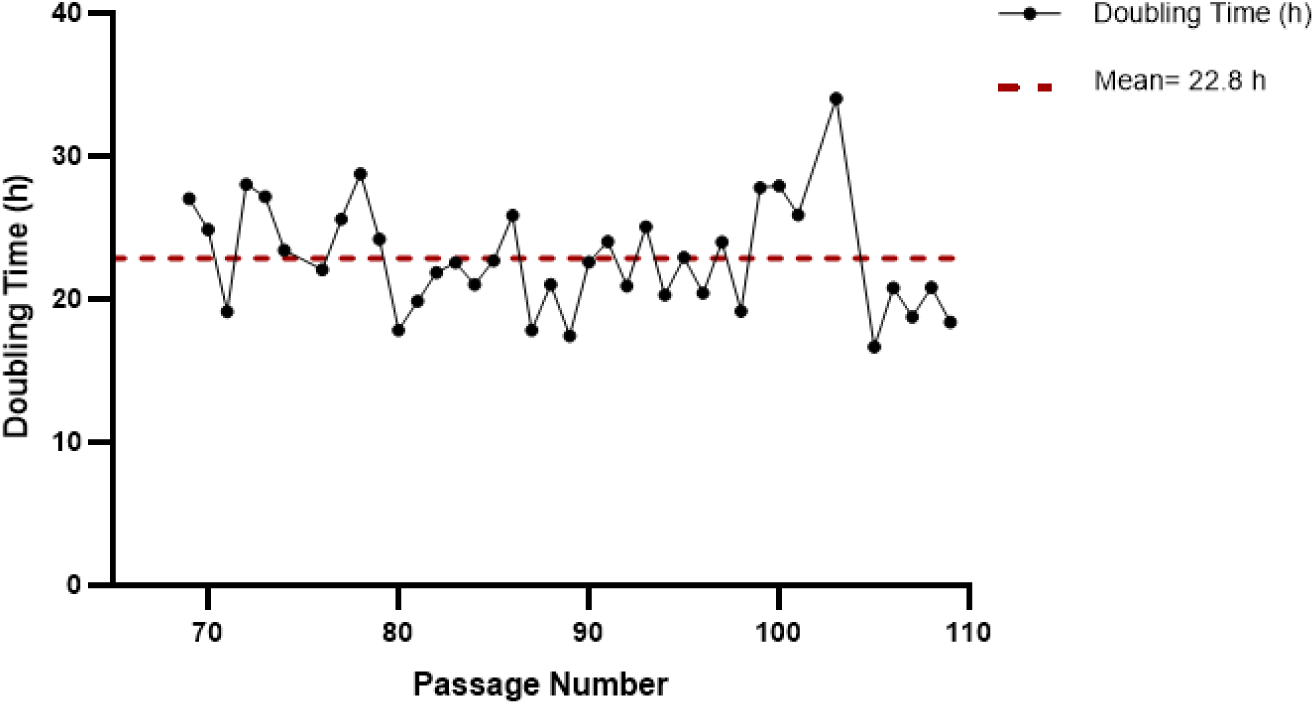
Population doubling time of cell line PE25 Working Cell Bank (WCB) cells across 40 passages. Doubling time was calculated from viable cell counts taken at defined time points during exponential growth at each passage interval. Mean doubling time of 22.8 h is indicated by the red dashed line. Each data point represents a single measurement (*n* = 1).

### p53-Mediated Apoptotic Response: PE25 WCB Cells Are Chemosensitive, Not Chemoresistant

Doxorubicin treatment produced markedly different dose-response profiles in PE25 WCB cells (passage 91) versus B16-F10 melanoma control cells (passage 6) (**Fig. 2 & 3**). At sub-lethal doxorubicin concentrations, B16-F10 cells exhibited an increase in proliferation rate, consistent with hormesis and documented as a survival adaptation in cancer cell biology^47^. This response was absent in PE25 WCB cells, which showed (**Fig. 2**) a monotonic decline in viability consistent with intact p53-mediated apoptotic machinery^48^. PE25 WCB cells exhibited a hypersensitive apoptotic response with an IC50 of 0.98±0.5 µM, indicative of a fully functional p53 pathway and intact genomic surveillance. B16-F10 cells demonstrated pronounced chemoresistance with an IC50 of 3.9±0.9 µM—a nearly four-fold difference (**Fig. 3**)—consistent with p53 pathway evasion in transformed cell lines^49,50,51^.

**Fig. 2.**
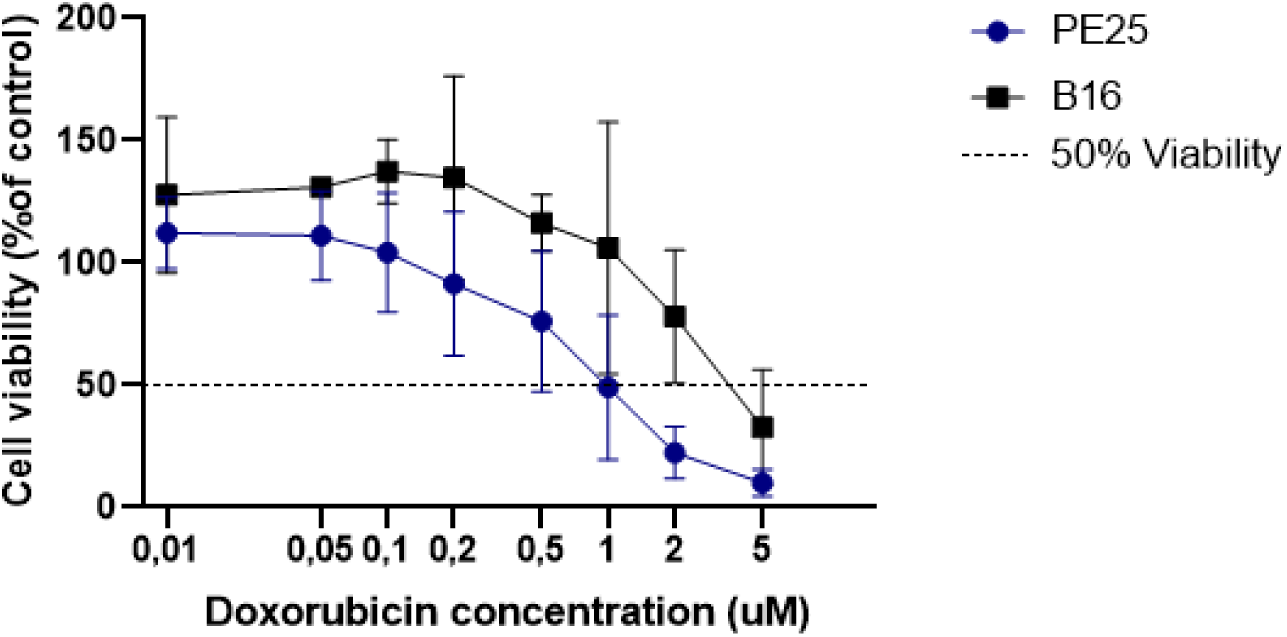
Dose response upon doxorubicin treatment. This graph shows the cell viability of both PE25 and B16 cell lines compared in % to the untreated control group (*n* = 3).

**Fig. 3.**
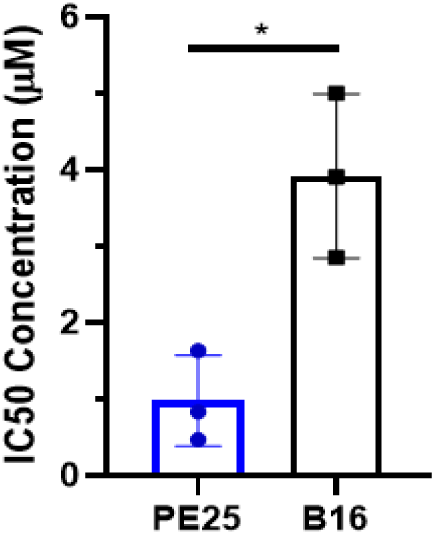
Comparative evaluation of half-maximal inhibitory concentrations (IC50) of doxorubicin across the stem cell line, PE25, and the malignant cell line, B16. Quantification of chemosensitivity thresholds in the mouse embryonic stem cell line, PE25, compared to the melanoma cell line, B16, following a 24-hour doxorubicin exposure. The non-transformed PE25 line demonstrates a highly sensitive, tightly regulated apoptotic threshold with a mean IC50 of 0.98±0.5 μM. In contrast, the transformed B16 cell line exhibits substantial chemoresistance, with an elevated mean threshold of 3.9±0.9 μM. Data are presented as mean ± standard deviation (SD) for biological triplicates, and statistical significance was determined using an unpaired t-test.

### Oncogenic Marker Expression: *CD44* and *BMI1* Are Not Upregulated in PE25 WCB Cells

qPCR quantification confirmed that PE25 WCB cells (passage 101) maintain transcriptional profiles for *CD44* and *BMI1* comparable to the non-transformed ES-E14TG2a mESC reference line (passage 14). Both markers were significantly upregulated in the B16-F10 melanoma cell line relative to PE25 and ES-E14TG2a (**Fig. 4**). The absence of *CD44* and *BMI1* upregulation is consistent with PE25 cells not having acquired the metastatic or dysregulated self-renewal signatures associated with spontaneous oncogenic transformation following extended passaging and serum-free adaptation^52,53,54^.

**Fig. 4.**
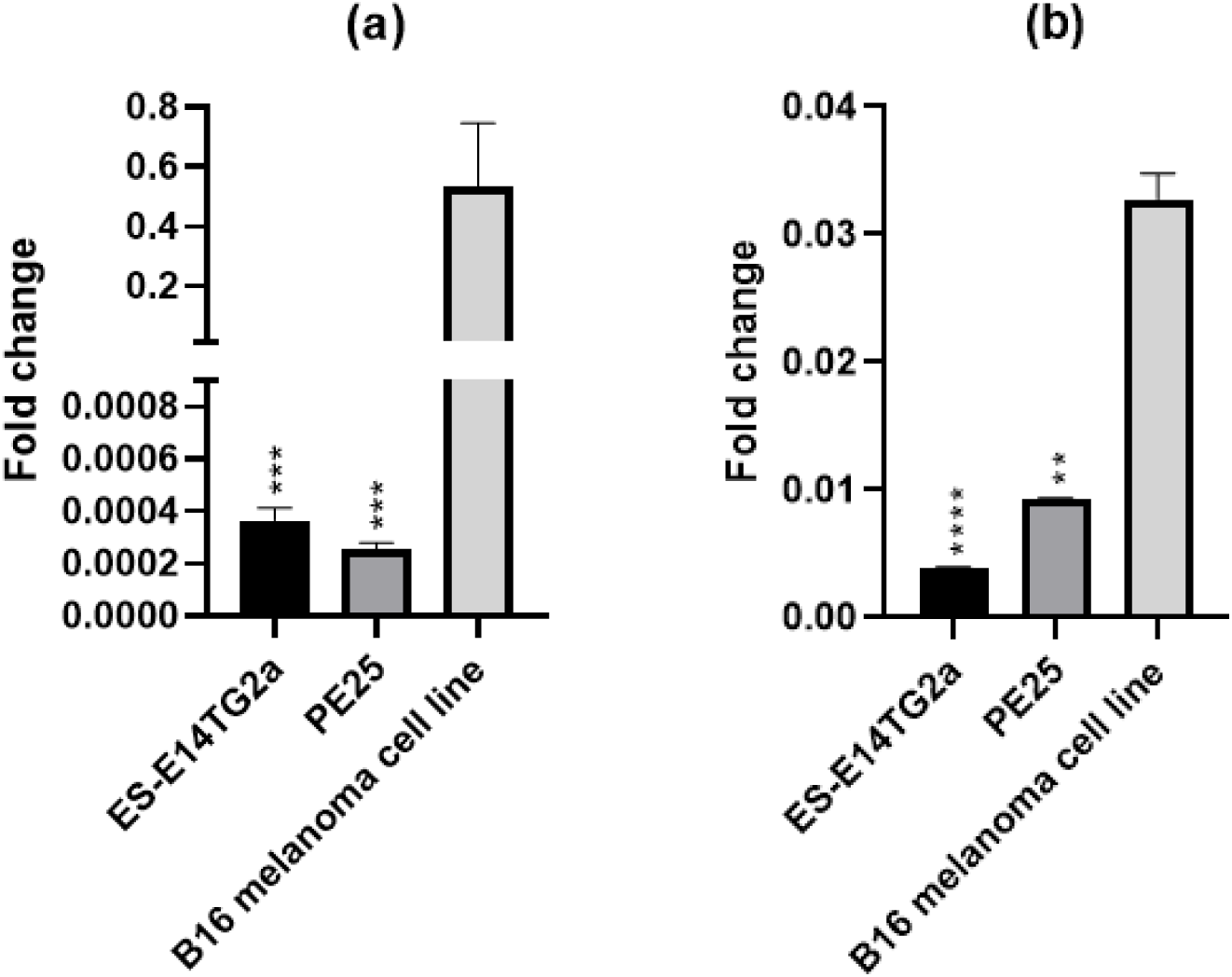
Relative mRNA expression levels of (a) *CD44* and (b) *BMI1* in PE25 compared to characterized ES-E14TG2a and B16 cell lines. The expression levels of *CD44* and *BMI1* in PE25 are comparable to the non-transformed ES-E14TG2a mESC reference line, indicating a lack of spontaneous oncogenic activation. Significant upregulation of both markers is observed in the B16 melanoma cell line, which are associated with increased metastatic potential and dysregulated self-renewal. Data are presented as fold change relative to housekeeping genes (*Actin* and *GAPDH*). Error bars represent the standard deviation (SD) of two biological replicates (*n* = 2). Statistical significance was determined using an ordinary one-way ANOVA with Dunnett’s multiple comparisons test, using the B16 melanoma cell line as the reference control.

### Soft Agar Assay: WCB Cells Do Not Exhibit Anchorage-Independent Growth

PE25 WCB cells (passage 94) remained entirely as single cells throughout and formed no colonies (**Fig. 5a**), demonstrating a complete absence of anchorage-independent growth—the hallmark phenotypic indicator of malignant transformation^55^. Conversely, B16-F10 melanoma cells (passage 6) formed robust multicellular spheroids and macroscopic colonies over the 17-day assay period (**Fig. 5b**).

**Fig. 5.**
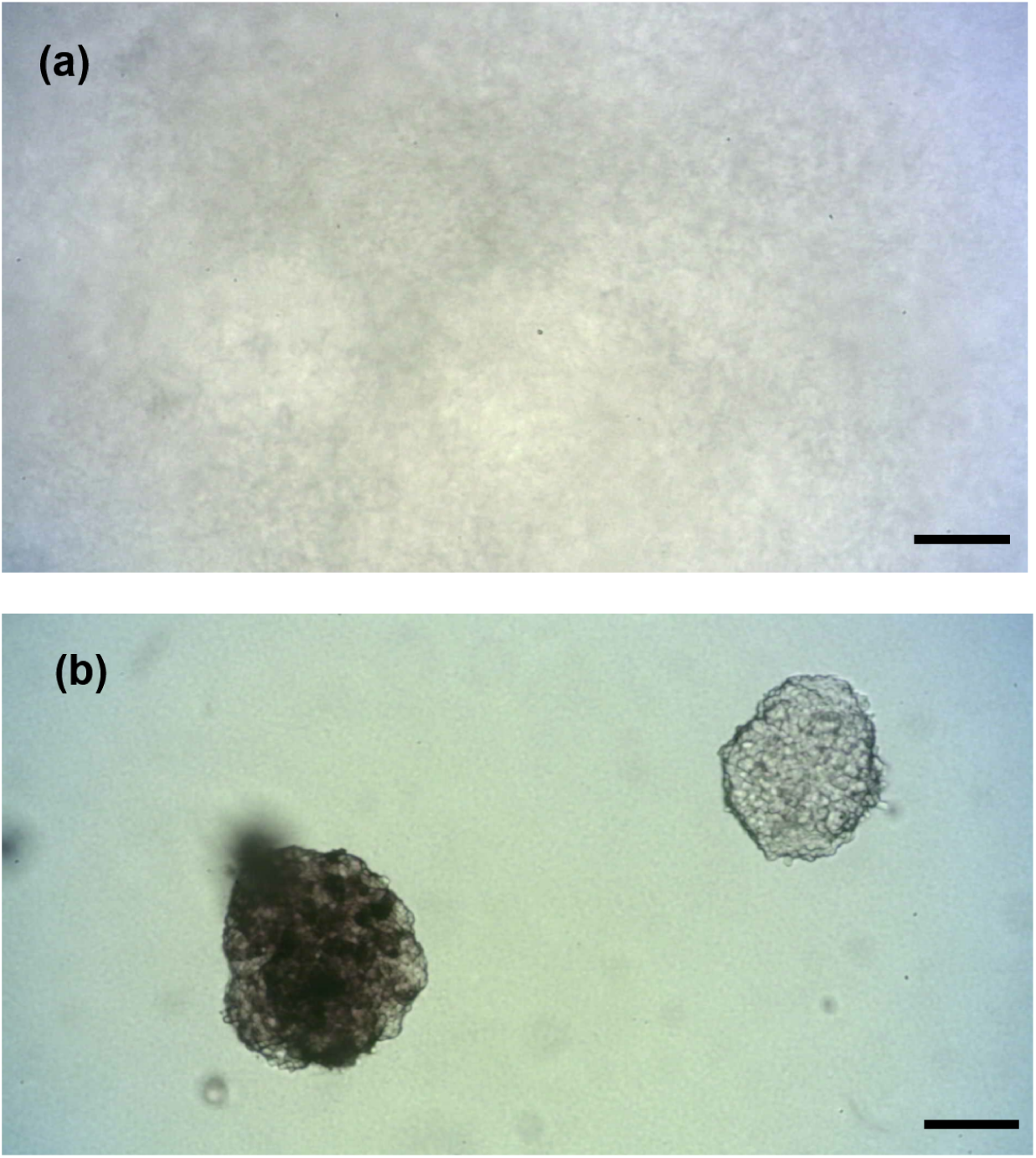
Soft agar colony formation assay comparing B16 mouse melanoma cells and PE25 WCB cells after 17 days. **(a)** PE25 WCB cells (passage 94) and **(b)** B16 mouse melanoma cells (passage 6), cancer-positive reference. Cells were suspended in 0.35% agarose over a 0.7% agarose base and incubated for 17 days. Images were acquired by brightfield microscopy at 5× magnification. Scale bar, 50 µm. The assay was performed in triplicate (*n* = 3).

Taken together with the p53 stress assay and oncogenic marker data, these results confirm that cell line PE25 is non-tumorigenic.

### Cell Viability Across Production Stages

CellTiter-Glo® measurements confirmed high viability in both the expansion bioreactor (∼98%) and the production bioreactor (∼85%). No viable cells were detected in the finished sample following the freeze-thaw cycle (**Fig. 6**). The final product—comprising both PE25 cell biomass and conditioned media—therefore consists exclusively of non-viable, inactivated cells with no detectable replicative capacity.

**Fig. 6.**
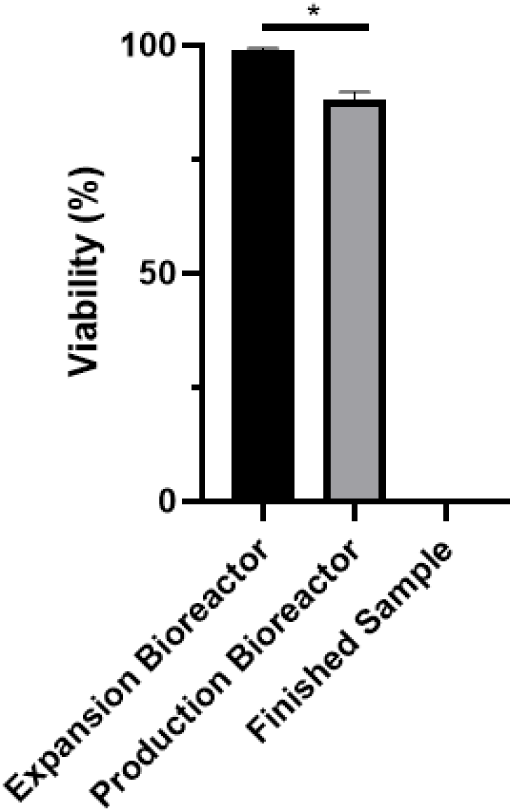
PE25 cell viability throughout production phases. Viability of cell line PE25 WCB cells was measured via CellTiter-Glo® assay (*n* = 3). High viability is maintained through the expansion and production bioreactor phases, while no viability is observed post-harvest following the freeze-thaw cycle. Data are presented as mean ± standard deviation (SD) for biological triplicates, and statistical significance was determined using a paired t-test.

### Genotoxicity: Ames Test and Micronucleus Assay

Third-party Ames testing conducted under OECD TG 471, using a treat-and-plate adaptation, was negative. No test item precipitation was observed at any concentration, and normal background lawn growth was maintained across all tester strains up to the maximum dose of 5,000 µg/plate, regardless of S9 metabolic activation status. Revertant colony counts remained above the 0.5 indication factor threshold, confirming the absence of cytotoxic effects. No significant increase in revertant numbers, nor any dose-dependent trend, was observed within the range of biological relevance. Positive controls produced the expected increases in revertant colonies within historical laboratory reference ranges, confirming assay validity. The ingredient therefore did not induce gene mutations via base pair substitutions or frameshifts under the reported conditions.

Independent third-party micronucleus testing under OECD TG 487 also returned a negative result. The ingredient’s solubility was confirmed in sterile water at 400 mg/mL, enabling evaluation at eight concentrations (0.0033, 0.0082, 0.0205, 0.0512, 0.128, 0.32, 0.8, and 2 mg/mL) alongside concurrent vehicle and positive controls across all three treatment setups. Cytotoxicity remained within acceptable regulatory limits (≤ 55 ± 5%) across all conditions, and concentrations from 0.32 to 2 mg/mL were selected for scoring. No statistically significant increase in micronuclei was observed at any tested concentration relative to the vehicle control. Positive controls demonstrated distinct, significant increases across all setups, fully validating the sensitivity and integrity of the test system. The PE25 cell line and conditioned media as an ingredient is therefore confirmed as neither clastogenic nor aneugenic.

## Discussion

### Cell Line PE25 Is Non-Tumorigenic: Convergent Evidence from Three Independent Assays

While genetic screening for p53 and Rb1 mutations is a traditional starting point for cell line safety assessment, genomic profiling alone is insufficient: it cannot capture epigenetic silencing of tumour suppressor loci, post-translational modifications that disable pathway components, or the emergence of subpopulations with malignant phenotypic capacity^56,57^. This study therefore combines functional and behavioral assays—specifically, the doxorubicin-induced stress assay and the soft agar colony formation assay—to directly evaluate p53 pathway integrity and anchorage-independent cell behavior. This functional approach was complemented by transcriptional profiling focused specifically on cancer stem cell (CSC) signatures (*CD44* and *BMI1*) rather than generic proliferation.

The doxorubicin stress assay demonstrates that the p53 tumour suppressor pathway remains fully functional in PE25 WCB cells following extended passaging and serum-free adaptation. The IC50 of 0.98 µM, the monotonic viability decline across the dose range, and the complete absence of sub-lethal proliferative hormesis collectively distinguish PE25 WCB cells from transformed lines in a quantitatively robust manner. The nearly four-fold difference in IC50 relative to B16-F10 melanoma cells, and the marked hormetic proliferative spike observed in B16-F10 but entirely absent in PE25, provide a clear contrast between a functional and a compromised p53 pathway.

The soft agar colony formation assay^58^ assesses anchorage-independent proliferation—a hallmark of malignant transformation—and evaluates tumorigenic potential directly through cellular behavior rather than through the identification of specific genomic or transcriptomic alterations^56,59^. PE25 WCB cells formed no colonies over 17 days and remained entirely as single cells, in direct contrast to the robust multicellular spheroids and macroscopic colonies formed by B16-F10 melanoma controls. This represents the gold-standard phenotypic test for malignant transformation: it is not a correlate or proxy, but a direct demonstration that PE25 cells lack the capacity for the hallmark behaviour of transformed lines.

The selection of *CD44* and *BMI1* as transcriptional endpoints was deliberate. Stem cells endogenously express high levels of self-renewal markers; generic proliferation markers therefore cannot discriminate a healthy stem cell from a transformed one in this context^60^. CD44, a cell-surface glycoprotein, and BMI1, a polycomb-group epigenetic regulator, are established master drivers of the CSC phenotype—dysregulated self-renewal, tumour initiation, and metastatic capacity—and critically, their upregulation is not a property of normal stem cells but specifically marks oncogenic transformation^61,62^. The absence of *CD44* and *BMI1* upregulation in PE25 WCB cells relative to the non-transformed ES-E14TG2a mESC control provides high-fidelity evidence that serum-free adaptation across extended passage has not driven acquisition of a CSC-like transcriptional programme. This baseline expression profile is further validated by the B16-F10 melanoma positive control line, which demonstrated the expected robust upregulation of both markers.

### Functional Genomic Stability in the Context of mESC Biology

Mouse embryonic stem cells are well-characterized in the literature as karyotypically dynamic: aneuploidy and chromosomal rearrangements accumulate readily during long-term culture, and this is an intrinsic biological property of the murine genome in pluripotent cells rather than an indicator of product-specific failure^63,64^. It would therefore be both scientifically inaccurate and misleading to claim karyotypic stability as a safety endpoint for a mESC-derived ingredient. The appropriate safety question is not whether chromosomal copy number variation is absent—it is unlikely to be—but whether any such variation has produced a population with compromised genomic surveillance, oncogenic transcriptional reprogramming, or capacity for anchorage-independent growth.

The data presented here address precisely these questions. Functional p53 pathway integrity confirms that cells sustaining consequential genomic damage are actively eliminated rather than selectively propagated. The absence of *CD44* and *BMI1* upregulation confirms that no oncogenic transcriptional programme has been activated. The negative soft agar result confirms that no subpopulation with malignant phenotypic capacity has emerged within detectable limits. Passage stability data—showing a consistent mean doubling time of 22.8 hours across 40 monitored passages with no systematic acceleration or deceleration—provides additional supporting evidence: erratic or progressively accelerating proliferative kinetics are a recognized correlate of genomic crisis, and their absence is consistent with a population that has reached a stable adapted state^11,17^. Taken together, this body of evidence supports a conclusion of functional genomic stability—the appropriate and scientifically defensible claim for an animal cell-cultured ingredient derived from mouse.

### Endogenous Retroviral Elements: Transparency Over Omission

The detection of MLV sequences in NGS transcriptomic analysis of the PE25 MCB is an expected and well-characterized feature of murine cell lines, reflecting the stable genomic integration of ERVs acquired through ancient germline infection events^40^. These sequences are present in essentially all *Mus musculus* cell lines and are distinguishable from exogenous, replication-competent retroviral contamination both by their integration pattern and by functional transmissibility assays. The confirmatory Q-PERT co-culture assay with *Mus dunni* target cells—a sensitive indicator cell line specifically chosen for its susceptibility to murine retroviruses—returned a negative result for replicative infectivity, directly demonstrating that no infectious viral particles capable of cell-to-cell transmission are produced. The decision to report these findings transparently, rather than omit NGS-detected ERV sequences from the public dossier, reflects the commitment to full disclosure motivating this work: a manufacturer that discloses negative outcomes on a sensitive viral screen demonstrates a more credible safety posture than one that reports no findings because the test was not performed.

### Genotoxicity Testing as a Core Safety Requirement for Animal Cell-Cultured Pet Food Ingredients

The Ames and in vitro micronucleus assays provide complementary coverage of the principal genotoxic hazards relevant to novel animal cell-cultured ingredients. The Ames test detects DNA-reactive mutagens that induce heritable gene mutations, while the micronucleus assay captures chromosomal damage, including both clastogenic (structural) and aneugenic (numerical) effects. Used together, these assays align with internationally accepted testing frameworks (e.g., OECD TG 471 and 487) and collectively address all three core genotoxic endpoints: gene mutation, structural chromosomal aberration, and numerical chromosomal aberration.

For animal cell-cultured ingredients, there is a strong rationale for testing the finished product rather than individual inputs. Production relies on complex media and generates diverse metabolic by-products; interactions within the harvested matrix may yield chemical species not detectable when components are assessed in isolation. Moveover studies confirm that extended cell culture produces metabolic profiles that diverge from primary cells and native tissues^65,66,67^, and can shift across passage within the same cell line, further increasing compositional variability^68,69,70^. This heterogeneity expands the space of potential molecular interactions, especially given that a substantial proportion of small molecules detected in untargeted metabolomics remains unannotated or only tentatively identified^71,72,73^.

Accordingly, cumulative genotoxic risk is most appropriately addressed through functional, whole-mixture testing. An in vitro screening battery combining the Ames test and the micronucleus assay provides coverage of both gene mutations and chromosomal damage in the final product^74,75^. For PE25 cells and conditioned media, negative results in both assays—across metabolic activation conditions and at concentrations up to 5,000 µg/plate in the Ames test and 2 mg/mL in the micronucleus assay—provide strong, independent evidence that the product does not pose a genotoxic hazard under the internationally accepted testing framework.

### Chemical Safety: DMSO Removal, Heavy Metals, Solvents, and Biogenic Amines

DMSO is universally employed as a cryoprotectant in cell banking at approximately 10% v/v and must be fully removed before the production seed train. The three step clearance protocol employed here—immediate 10-fold dilution at thaw, double wash, followed by a minimum of eight passages of 1:5 serial dilution in DMSO free media—is grounded in well characterized physicochemical principles governing passive membrane diffusion and concentration gradient driven efflux^41,42,43^. Third-party quantification at passage 8 confirmed DMSO below the limit of quantification (1 mg/kg), and conservative worst case modelling places residual DMSO exposure in a consuming animal at greater than 88,000-fold below the established chronic toxicity threshold in dogs^44^. This analysis not only confirms safety but provides the industry with a transparent methodological framework for DMSO clearance documentation.

Although no solvents other than DMSO—used solely at the cell banking stage and subsequently removed—were employed in this production process, screening for a comprehensive panel of organic solvent residuals remains important and was conducted here. Even where solvents are not directly used in manufacturing, food-grade and feed-grade ingredients—including amino acids, vitamins, and other media components—can introduce trace solvent residuals into a product, as solvents are routinely employed in the synthesis, extraction, and purification of these materials and may persist at low levels despite removal steps^37,76^. Testing the finished ingredient for solvent residuals is therefore an independent and necessary safety verification step, regardless of whether solvents were deliberately used in production. The detected ethanol level of 6mg/kg (LOQ 1mg/kg) in the product analyzed here represents an unintentional and unavoidable technical residue compliant with Directive 2009/32/EC frameworks. Moreover, this minute trace sits >800-fold lower than the 5000 mg/kg maximum limit recommended for class 3 low toxicity solvents under VICH GL18 guidelines, confirming a negligible toxicological profile and full regulatory alignment^77^.

All other measured volatile organics solvents, heavy metals (Pb, Cd, As, Hg), BME and biogenic amines of potential concern fell below applicable limits of detection. In the case of spermine, while 70 mg/kg slightly exceeds the LOQ, it sits well below the 100 to 150+ mg/kg concentrations routinely found in conventional organ meats such as liver and thymus commonly consumed by companion animals^78^. Furthermore, spermine is a biogenic amine that is increasingly being investigated for its functional health benefits and is not considered a metabolite of concern^79,80^.

### Media Transparency: A Prerequisite for Independent Safety Assessment

The full disclosure of production media components and their regulatory status is a prerequisite for meaningful independent safety assessment. Without knowledge of what the animal cell-cultured ingredient was grown in, neither regulators nor downstream manufacturers can assess contamination risk plausibility, evaluate residue screening completeness, or make informed decisions about ingredient suitability. The confirmation that all production media components—spanning amino acids, vitamins, minerals, metabolic compounds, protein and antioxidants—hold current EU Feed Additive approval status or are listed in the Feed Material catalogue establishes a clear regulatory baseline and eliminates uncertainty about the chemical provenance of the final product.

### Cell Viability in the Final Product

Where an animal cell-cultured ingredient for pet food is registered under the EU Feed Materials framework rather than subject to Novel Food authorization, no specific characterization requirements for cell viability are mandated. Nonetheless, EFSA’s 2024 updated Guidance on the scientific requirements for novel food applications under Regulation (EU) 2015/2283 provides a useful and well considered reference point for ingredient-level characterization of animal cell-cultured products^31^. That guidance identifies the respective concentrations of viable and non-viable cells in a novel food biomass as a relevant characterization parameter that should be reported—a principle that translates reasonably to any cell-cultured ingredient regardless of the regulatory route taken. It is worth noting that this parameter reflects the importance of clearly defining what a novel ingredient is, rather than implying that viable cells represent a safety hazard per se. The scientific consensus holds that ingested cells—whether viable or not—are broken down by the digestive system into their constituent molecular components and do not present a systemic risk to the consuming animal; this is equally true of the many viable cells naturally present in conventional whole foods. CellTiter-Glo® measurements confirmed that the freeze-thaw cycle employed in the production process results in complete cell inactivation: no viable cells were detectable in the final harvested ingredient. The final product is therefore a stable matrix of non-viable cell biomass and conditioned media—a characterization considered relevant both for alignment with best-practice scientific standards and for the confidence of consumers and downstream manufacturers who reasonably wish to understand what the ingredient contains.

### Each Cell-Cultured Ingredient Must Be Evaluated Individually

The characterization data presented here apply specifically to cell line PE25 and its derived ingredient and cannot be extrapolated to other animal cell-cultured ingredients. Cell line PE25 was derived from a non-genetically modified embryonic stem cell background and adapted to food-safe, feed-grade media. This represents a fundamentally different biological starting point from deliberately immortalized lines or spontaneously transformed/adapted somatic cell lines^7,8,9^. The functional integrity of the p53 pathway demonstrated here directly addresses a common molecular basis of cancer-like phenotypes in cultured animal cell lines^17^; and its preservation following extended passaging in serum-free, food-safe, feed-grade media represents a meaningful safety distinction. Whether cell lines produced by other manufacturers can make equivalent claims is a question that only publicly disclosed, product-specific characterization data can answer^11^.

### Voluntary Transparency as an Ethical and Commercial Standard

The current EU and UK regulatory frameworks permit animal cell-cultured pet food ingredients to reach the market without independent ingredient-level safety review. In this context, voluntary public disclosure of comprehensive safety data is simultaneously an ethical obligation, a prerequisite for downstream supply chain confidence, and the most durable foundation for long-term consumer trust. This paper represents the first public disclosure of a comprehensive safety dossier for any such ingredient. The proposed minimum disclosure framework presented here is not exhaustive, but it defines the core evidentiary suite that any responsible claim of commercial readiness should be prepared to support publicly and permanently.

Within this framework, we show that cell line PE25 is non-tumorigenic and functionally genomically stable, and that its derived ingredient is non-mutagenic, non-clastogenic, free of heavy metal and solvent exceedances, and is produced using components that hold current approval as EU Feed Additives or are listed in the EU Feed Materials catalogue. These data illustrate how a full safety dossier can be made transparently available without regulatory compulsion and underscore that each animal cell-cultured ingredient must be evaluated on its own merits. We commit to this standard and call on the broader industry to do the same.

## Materials and Methods

### PE25 Cell Line Origin and Derivation

Mouse embryonic stem cells (*Mus musculus*) were isolated from the inner cell mass of blastocysts and plated under sterile, serum-free, feeder-free conditions. Clones displaying canonical mESC morphology were expanded under established 2D adherent protocols with daily media exchange and enzymatic passaging every 3–4 days. A Parental Cell Bank (PCB) for each embryonic stem cell line was generated at passage 3 or earlier.

### Commercial Cell Lines

The mouse embryonic stem cell line ES-E14TG2a (ATCC® CRL-1821™; RRID:CVCL_9108) was obtained from the American Type Culture Collection (ATCC, Manassas, VA, USA) and used as a non-transformed mESC reference. The B16-F10 mouse melanoma cell line (ATCC® CRL-6475 ™) was also obtained from the American Type Culture Collection (ATCC, Manassas, VA, USA) and used as a cancer-positive reference control.

### Creation of the PE25 Master Cell Bank (MCB)

The MCB for the mouse embryonic stem cell line PE25 was established from the PE25 PCB following ICH Q5D guidelines from a single pooled culture to ensure homogeneity across all cryovials. Cells were expanded through 250 mL 3D suspension cultures and banked at passage 16. A total of 95 cryovials were produced in DMSO-free Bambanker Cell Freezing Medium, slow-frozen at -80°C for 24 hours using a Corning CoolCell container (Corning, USA), and transferred to liquid nitrogen dewars for long-term storage. Independent safety validation of the MCB was conducted by Clean Cells SAS (Montaigu-Vendée, France) and PathoQuest SAS (Paris, France). Species-specific identity and cross-contamination screens were performed via qPCR targeting *Mus musculus* genomic markers. Sterility testing was executed under good manufacturing practices (GMP) via direct inoculation methods. The assay evaluated potential contamination using a panel of validated positive control strains and targeted qPCR assays were performed to screen for mycoplasma and spiroplasma contamination according to EP 2.6.7 and USP 63 guidelines. Broad-spectrum adventitious virus detection was conducted using NGS transcriptomic profiling. To address the regulatory significance of endogenous MLV sequences identified during transcriptomic mapping, Quantitative Product Enhanced Reverse Transcriptase (Q-PERT) assay was performed. MCB test cells were co-cultured with target *Mus dunni* cells to monitor viral transmissibility, utilizing a highly sensitive product-enhanced reverse transcriptase (Q-PERT) readout.

### Creation of the PE25 Working Cell Bank (WCB)

The WCB was created from the PE25 MCB following ICH Q5D guidelines. Cells were progressively weaned from research-grade to food-safe and feed-grade media and passaged 46 times across multiple bioreactors. All components of the food-safe and feed-grade media are either approved EU Feed Additives or listed as EU Feed Material (**Table 1**). A WCB of a total of 20 cryogenic vials (Corning, USA) was produced, each containing 1 mL of cell suspension in food-safe and feed-grade media supplemented with 10% DMSO as a cryoprotectant.

From the initial cell line derivation through the establishment of the MCB and WCB, all handling, documentation, and traceability were carried out in strict compliance with Regulation (EC) No 1069/2009 on animal by-products not intended for human consumption^81^.

### Heavy Metal Screening

Biomass and conditioned media from PE25 WCB expansion and scale-up were submitted to SGS Germany GmbH (Hamburg, Germany)—a Deutsche Akkreditierungsstelle (DAkkS)-accredited laboratory with DIN EN ISO/IEC 17025 accreditation—for analysis of lead (Pb), cadmium (Cd), arsenic (As), and mercury (Hg) by ICP-MS in accordance with DIN EN 15763, mod. Results were evaluated against the maximum permitted levels established in EU Directive 2002/32/EC.

### Solvent and Chemical Reagent Residue Analysis

Biomass and conditioned media from PE25 WCB expansion and scale-up was screened for the solvents benzene, methanol, ethanol, acetone, isopropanol, toluene, ethylbenzene, m,p-xylene, o-xylene, chloroform, dichloromethane, trichloroethylene, 1,1,1-thrichloroethane, tetrachloroethylene and carbon tetrachloride by headspace gas chromatography–mass spectrometry (HS-GC/MS) in accordance with SOP M 1299 (SGS Germany GmbH, Hamburg, Germany). DMSO was assayed by PiCA Prüfinstitut Chemische Analytik GmbH (Berlin, Germany), a DAkkS-accredited laboratory, using GC-MS. BME was assayed by Analytice (Strasbourg, France), operating under ISO 9001 and 14001 quality management systems, using GC-MS. In the absence of a feed-specific residual solvent standard, results were benchmarked against the limits established for human food, pharma and veterinary products. All results were evaluated against the permitted daily exposure limits and purity requirements of EU directive 2009/32/EC (extraction solvents in food), ICH Q3C (residual solvents scientific guideline) and VICH GL18 (residual solvents in veterinary medicinal products). To ensure comprehensive compliance with the feed safety criteria, results were further cross-referenced with applicable EU feed safety standards, specially regulation (EU) No 68/2013 (technical processing residues), and regulation (EC) No 183/2005^82^ (feed hygiene).

### Biogenic Amine Analysis

Histamine, cadaverine, putrescine, spermidine, spermine, tyramine, phenylethylamine, tryptamine, and isopentylamine were assayed from PE25 WCB-generated cells and conditioned media by SGS Germany GmbH (Hamburg, Germany) using reversed-phase high-performance liquid chromatography (RP-HPLC, SOP M 1574).

### Passage Stability: Doubling Time Monitoring

To assess proliferative stability of PE25 WCB cells across the production passage range, cell doubling time was measured at regular passage intervals throughout the expansion process. Cell counts were performed at each passage using high-precision automated cell counter NucleoCounter^®^ NC-200™ (ChemoMetec, Denmark), and doubling time was calculated from the formula td = t / [(log Nt − log N0) / 0.301], where *t* is elapsed culture time, *N*0 is the total inoculation number, and *N*t is the total harvest number. Data are presented as doubling time (hours) as a function of passage number. All measurements were taken from routine production monitoring records (*n* = 1 per passage); this was not a dedicated replicated experiment.

### Doxorubicin-Induced Stress Assay (p53 Pathway Integrity)

To assess functional p53 pathway integrity—a critical safety indicator for evaluating intact tumor suppressor mechanisms and cellular response to DNA damage—PE25 WCB cells (passage 91) and B16-F10 melanoma cells (passage 6) were seeded at equivalent densities and treated with a serial dilution of doxorubicin hydrochloride for 24 hours. Cell viability was measured using the CellTiter-Glo® Luminescent Cell Viability Assay (Promega, USA). All conditions were performed in biological triplicate (*n* = 3). Half-maximal inhibitory concentration (IC50) values were calculated from dose-response curves, and viability data were normalized to untreated controls (100%).

### Transcriptional Profiling of Oncogenic Markers (*CD44* and *BMI1*) of PE25 WCB cells

To characterize the stemness, self-renewal capacity, and potential oncogenic phenotype of the cell line, mRNA expression of the cancer stem cell markers *CD44* and *BMI1* was evaluated. Total RNA was extracted from PE25 WCB cells (passage 101), B16-F10 mouse melanoma cells (cancer-positive reference, passage 6), and ES-E14TG2a (non-transformed negative reference, passage 14) via phenol/chloroform extraction^83^. cDNA was synthesized and quantitative polymerase chain reaction (qPCR) was performed to quantify mRNA expression of *CD44* and *BMI1*. Data are presented as fold change relative to housekeeping genes *Actin* and *GAPDH*. All measurements were performed in biological duplicates (*n* = 2); error bars represent standard deviation (SD).

### Soft Agar Colony Formation Assay

Anchorage-independent growth—the ability of cells to proliferate without adhering to a solid substrate—was assessed using a standard soft agar colony formation assay. Historically recognized in the pharmaceutical industry as the gold standard *in vitro* assay for validating cellular transformation, this method serves as a critical benchmark for evaluating tumorigenic potential and phenotypic stability. To evaluate this trait, the soft agar assay was performed with slight adaptations^55^. PE25 WCB cells (passage 94) and B16-F10 mouse melanoma cells (passage 6) were suspended in 0.35% agarose layered over a 0.7% agarose base in complete medium and incubated for 17 days. Colonies and cell morphology were visualized by brightfield microscopy at 5× magnification. The assay was performed in triplicate (*n* = 3).

### Cell Viability Across Production Stages

To evaluate the physiological health and phenotypic stability of the cell line across the manufacturing workflow, cell viability was assessed at three sequential production stages: the expansion bioreactor, the production bioreactor, and the post-harvest final sample (following freeze-thaw cycle). Monitoring cell viability is a critical component of culture characterization, as a significant drop in viability serves as a direct indicator of a failing culture or adverse processing conditions. All measurements were performed in triplicate (*n* = 3) using the CellTiter-Glo® Luminescent Cell Viability Assay, and with data normalized to the expansion bioreactor stage as 100% baseline.

### Ames Test (Bacterial Reverse Mutation Assay)

Third-party mutagenicity testing of PE25 WCB-generated cells (passage 82) and conditioned media was performed by GBA ICCR-Roßdorf GmbH (Roßdorf, Germany), a DAkkS-accredited laboratory, using the Ames bacterial reverse mutation assay (OECD Test Guideline 471) under good laboratory practice (GLP) conditions^32^. The study employed the treat-and-plate method with *Salmonella typhimurium* tester strains TA 98, TA 100, TA 102, TA 1535, and TA 1537, with and without metabolic activation (S9 fraction), in triplicate (*n* = 3). The evaluation was conducted in two phases: Phase I (pre-experiment) at dose concentrations of 100, 333, 1000, 2500, and 5000 µg/plate, and Phase II at 312.5, 625, 1250, 2500, and 5000 µg/plate.

### Micronucleus Assay (Mammalian Cell Genotoxicity)

Third-party chromosomal genotoxicity testing of PE25 WCB-generated cells (passage 82) and conditioned media was performed by Syngene International Ltd (Bangalore, India), accredited to ISO/IEC 17025:2017, ISO 15189:2012, and ISO 9001:2015, using the in vitro mammalian cell micronucleus test^33^ (OECD Test Guideline 487). Human peripheral blood lymphocytes were exposed to the test item across three treatment setups: 4-hour treatments with and without rat liver S9 metabolic activation, and a 24-hour treatment without S9, each in duplicate (*n* = 2). Following cytochalasin B block, 2,000 binucleated cells per group were scored for micronucleus frequency and cytotoxicity. The assay assessed both clastogenic and aneugenic potential.

### Media Component Regulatory Assessment

All components of the production media were assessed for regulatory approval status under EU feed legislation by cross-referencing each ingredient against the EU Register of Feed Materials (Regulation (EU) No 68/2013) and the EU Register of Feed Additives (Regulation (EC) No 1831/2003). Approval status, regulatory category, and the applicable register entry for each component are presented in **Table 1**.

## Statistical analysis

Unless otherwise specified, a two-way analysis of variance (ANOVA) was conducted to facilitate comparison between multiple groups and pairwise groups. The outcomes of the experiments were visually represented using GraphPad Prism version 8.0.0 (GraphPad Software, La Jolla, CA, USA). Quantitative data are presented as mean values accompanied by their respective standard deviations (SD) for biological triplicates (*n* = 3). Long-term passage kinetic tracking (**Fig. 1**) represents single sequential run metrics (*n* = 1 per passage interval) extracted from real-time production logs. Statistical significance was defined as p < 0.05. Probability values are represented in figures as follows: *p < 0.05, **p < 0.01, ***p < 0.001, and ****p < 0.0001.

## Data availability

The data that support the safety conclusions of this study are available within this article. Third-party test certificates supporting the reported results are available from the corresponding author upon reasonable request.

## Acknowledgements

The authors thank Dr. Dima Faour-Klingbeil for kindly reviewing the manuscript and providing valuable insights. This work was supported by the Austrian Research Promotion Agency (Österreichische Forschungsförderungsgesellschaft, FFG).

## Author contributions

R.T., R.S. and S.F. conceptualized the study. R.S., M.S., M.F., M.M. and L.H. performed the in-house assays and analyzed the data. S.F. provided overall supervision and acquired funding. R.T., R.S. and S.F. prepared the original manuscript draft and contributed to the review and editing of the manuscript.

